# The marine microbiome can accurately predict its chemical and biological environment

**DOI:** 10.64898/2026.01.25.701552

**Authors:** Emma Bell, Karin Garefelt, Krzysztof T Jurdzinski, Luis Fernando Delgado, Fanny Johansson, Bengt Karlson, Anders F. Andersson

## Abstract

The microbiome responds to physicochemical changes in the environment, making it a sensitive indicator of ecosystem status. Monitoring microbial communities in aquatic systems is therefore essential for understanding ecosystem health and responses to change. Traditionally reliant on microscopy, monitoring programmes are increasingly incorporating DNA-based approaches leveraging on advances in high-throughput sequencing. In this study, we evaluate the potential of using DNA metabarcoding to predict abiotic and biotic parameters across the spatiotemporal gradients of the Baltic Sea. The dataset comprises 397 seawater samples integrating prokaryotic (16S rRNA gene) and eukaryotic (18S rRNA gene) metabarcoding data with environmental measurements and plankton microscopy counts. Random Forest models based on metabarcoding data accurately predicted a range of physicochemical parameters and showed performance comparably to more complex machine learning algorithms. Models based on 16S rRNA gene data tended to perform better than those based on 18S rRNA gene data, with amplicon sequence variant-level data yielding the best results. Metabarcoding outperformed plankton microscopy in predicting abiotic factors and effectively predicted the presence of phytoplankton and zooplankton genera using ≤1 L of water. Models trained on independent datasets accurately predicted several of the physicochemical parameters, but performed weaker on others, highlighting the potential and challenges for their transferability. Furthermore, our predictions closely matched the observed HELCOM indicator values for assessing good environmental status, suggesting the utility of microbiome-based approaches in regional marine monitoring frameworks. These findings underscore the potential of environmental DNA as a tool for ecosystem monitoring and management in dynamic coastal systems.

## Introduction

Microorganisms and the ecological niches they inhabit have a reciprocal relationship. Microbial metabolism drives element and nutrient cycles and is constrained by environmental factors such as temperature, pH, salinity, redox potential, and available carbon and nutrient sources. At the same time, the physicochemical environment shapes the composition of the microbiome, since each microbial strain has a unique fitness distribution in a multidimensional niche space [1, 2]. The close link between microorganisms and their ecological niche means microbiome data can provide insight into the status of the ecosystem [3–6].

Marine environmental monitoring typically includes measurements of physical, chemical, and biological parameters. Microscopy is the traditional method for monitoring marine microorganisms and has been used for decades in long-term monitoring programmes [7–9]. The morpho-taxonomic data provides crucial information on the ecology of key plankton taxa and can serve as an indicator of ecosystem state [10]. However, the method is time-consuming, requires taxonomic expertise, and has limited taxonomic scope. Importantly, small microbes such as picoeukaryotes and bacterioplankton cannot typically be identified taxonomically despite carrying out important ecosystem functions. In contrast, metabarcoding and shotgun metagenomics make it possible to study entire microbial communities directly from environmental DNA [11, 12]. With the decreasing cost of high-throughput sequencing, these methods have become widely accessible, facilitating large-scale studies of marine microbial diversity [13], and are increasingly argued to be included in routine monitoring programmes [14, 15].

Recent research has adopted machine learning methods, such as Random Forests or Support Vector Machines, to harness the predictive power of microbiome data to categorise or forecast ecosystem conditions [16, 17]. Metabarcoding and metagenomics data have, for example, been used to predict soil health [4], detect contamination in groundwater and sediment [18–20], predict physicochemical properties of lake water [21], dissolved organic carbon in soil [22], and environmental status of freshwater reservoirs [23] and headwater streams [24]. While most studies employing machine learning for this purpose train the models directly on the sequence counts, sometimes aggregated at different taxonomic levels, recent studies have also employed deep learning approaches to obtain a smaller set of informative features to train the models on [25]. A key challenge in generating accurate predictions is obtaining high-quality training datasets that capture the full range of an ecosystem’s variability, including environmental gradients [26, 27], seasonal changes, and response to disturbances [28], which are needed to reliably train machine learning models [29].

Machine learning using metabarcoding data has been used in a marine context, for example for assessing the environmental impact of marine aquaculture [30–32] or predicting bacterial production [33, 34], but a systematic assessment of how effectively the microbiome can predict different environmental parameters of seawater is missing. Moreover, since many monitoring programs involve microscopy-based plankton identification, a comparison of the predictive power of the morpho-taxonomic data and metabarcoding data is relevant. Similarly, predicting microscopy counts would make metabarcoding studies more comparable with long-term microscopy-based time series, but has not been attempted yet. Finally, the transferability of microbiome-based predictive models when applied to a different dataset remains to be assessed.

Here, we use metabarcoding data to predict abiotic (physicochemical) and biotic (phytoplankton and zooplankton composition) parameters in the Baltic Sea (Fig. 1). We used a high-quality spatiotemporal dataset of prokaryotic and eukaryotic microbial communities that incorporates physicochemical and metabarcoding data from 397 samples sampled at 18 stations across the Baltic Sea, Kattegat, and Skagerrak [35, 36]. Changes in time (seasonality) and space (geographical region and salinity gradient) are reflected in the data, making it ideal for assessing the feasibility of using metabarcoding data as an environmental biosensor. We systematically evaluate the prediction performance of prokaryotic and eukaryotic marker genes (16S rRNA and 18S rRNA) at different taxonomic levels, feature extraction techniques, machine learning algorithms, and compare the performance of microscopy-based and metabarcoding-based predictions. Our dataset comprised two separate sampling and sequencing efforts, which allowed us to compare interannual predictions across different datasets with training and deploying on the same dataset. We further demonstrate this potential by predicting Baltic Marine Environment Protection Commission (HELCOM) indicators [37], which inform policy and management decisions to protect the ecological status of the Baltic Sea.

**Fig. 1.**
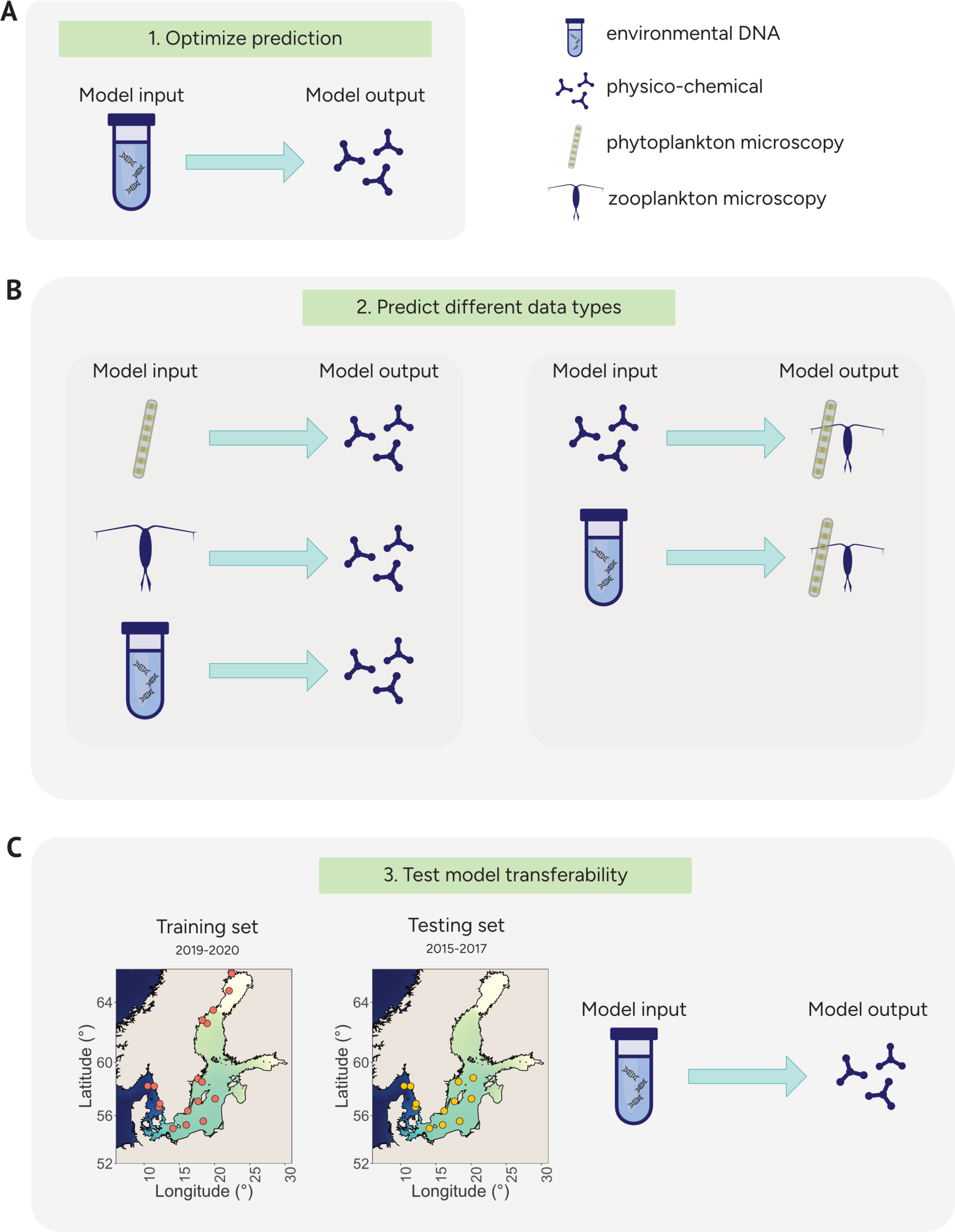
Overview of study design. **(A)** Evaluation of different machine learning architectures and feature extraction techniques for physicochemical data prediction from metabarcoding data. **(B)** Comparison of physiochemical predictions based on metabarcoding and microscopy data, and of microscopy plankton data using metabarcoding or physiochemical data. **(C)** Evaluation of model performance across datasets.

## Materials and Methods

### Datasets

Environmental data from the Swedish pelagic marine monitoring program is collected according to the guidelines of the HELCOM-COMBINE manual [38]. The data is freely available, and was downloaded from the SHARK database: https://shark.smhi.se/hamta-data/ and includes a range of physical, chemical, and biological properties of seawater. We selected physicochemical parameters measured for at least 10% of the samples in the dataset: salinity, temperature, pH, alkalinity, Secchi depth, SiO_3_, N_tot_, dissolved inorganic nitrogen (DIN), NH_4+_, NO_2-_, NO_3-_, combined measurements of NO_3-_ and NO_2-_, P_tot_, PO_4_, dissolved organic carbon (DOC), humus, and chlorophyll-a. In addition, sample date and time, longitude, and latitude are available for all samples. In total, 397 samples were used to create models from 16S rRNA or 18S rRNA gene amplicon data, with one sample from the original dataset [36] missing physicochemical data. The average data input for each parameter was 332 data points ranging from *n* = 75 (DOC) to *n* = 378 (temperature) (Supplementary Fig. 1).

Microscopy-based counts of phytoplankton and zooplankton from the Swedish pelagic marine monitoring program were also accessed from the SHARK database. Phytoplankton were sampled using an integrated hose sample, typically from 0–10 m depth, preserved in Lugol’s solution, settled in sedimentation chambers, and identified and counted with inverted light microscopy [39] and reported as individuals per L. Zooplankton were collected with vertically towed nets, with depth ranges varying between stations (from 0–6 m to 0–193 m depth). The zooplankton were identified and counted with light microscopy, and abundances were reported as individuals per m³. For each taxon, we summed counts across size classes (phytoplankton) and growth stages and sexes (zooplankton) when applicable. Only taxa identified to at least the genus level, and for zooplankton belonging to Arthropoda and Rotifera, were included in the analyses. Samples with both metabarcoding and plankton data were available from 18 stations for phytoplankton and 16 stations for zooplankton. When predicting physicochemical parameters using microscopy data, plankton counts (individuals per volume unit) were used. When predicting microscopy identifications of plankton genera using metabarcoding or physicochemical data, microscopy counts were first aggregated at the genus level, and then either individuals per volume unit (Supplementary Fig. 2) or presence/absence (main results) were predicted. For the latter, the microscopy count data was converted to binary data (presence/absence).

Biomass for microbial DNA has been collected and sequenced using DNA metabarcoding in parallel with the pelagic marine monitoring program in multiple sampling efforts. We used amplicon sequencing data previously published with ENA accession numbers PRJEB55296 [35] and PRJEB84926 [36]. The amplicon datasets contain both 16S rRNA (V3-V4 region) and 18S rRNA (V4 region) gene metabarcoding that capture prokaryotic and eukaryotic communities, respectively. Amplicon datasets were processed with the nf-core pipeline ampliseq [40], which implements DADA2 [41] to produce amplicon sequence variants (ASVs). ASVs were taxonomically annotated within DADA2 using a curated version [42] of the Genome Taxonomy Database [43] for 16S rRNA gene and the Protist Ribosomal Reference database [44] for 18S rRNA gene. For 16S rRNA gene, ASVs were also annotated with SILVA [45] (version 138.1) in order to identify chloroplast and mitochondrial sequences. Further details on the amplicon data processing are available in Latz et al. (2024) [35] and Jurdzinski et al. (2024) [36].

### Preparing the input data (features)

ASVs annotated as metazoa (18S rRNA) and chloroplast and mitochondria (16S rRNA) were removed from the dataset. Low prevalence ASVs that were present in ≤10% of all samples were excluded from the count tables to limit the number of features. Feature tables at higher taxonomic classification were created from aggregated ASV counts at the taxonomic levels: phylum, class, order, family, genus, and species for the 16S rRNA gene, and supergroup, division, subdivision, class, order, family, genus, and species for the 18S rRNA gene. Count tables were normalised by dividing raw counts by the total counts (excluding removed features) of each sample.

Deep encodings of the filtered and normalised ASV count data were created using autoencoders from a deep representation learning framework DeepMicro [25]. Multiple encoders from the DeepMicro framework were used: Shallow Autoencoders (SAE), Deep Autoencoders (DAE), Variational Autoencoders (VAE), and Convolutional Autoencoders (CAE). All configurations listed in Table S1 of the DeepMicro study [25] were used as dimensions for the networks.

Microscopy counts of zooplankton and phytoplankton were also used as features. Features were limited to taxa present in >10% of the samples for training the machine learning models. Taxonomic resolution was generally at the genus level, and in some cases species-level annotations were available.

### Predicting physicochemical properties

We trained three model architectures, Random Forest, XGBoost, and TabPFN, to predict physicochemical parameters of seawater in addition to contextual data. Data from all years (2015–2019) were included using 10-fold cross-validation. The folds for cross-validation were determined using ‘vfold_cv’ in the package rsample [46] with splitting based on the response variable.

Random Forest models were made with the R package ranger using default settings except for the number of trees, which was set to 2000. XGBoost models were trained using nested cross-validation with the R package caret (v7.0.1) [47]. For each parameter (response variable), the data was split into 10 outer folds. Within each training outer fold, hyperparameter tuning was performed using 10-fold inner cross-validation. A grid search was conducted across a set of hyperparameters with varying nrounds (200-1000 in steps of 50), eta (0.005, 0.010, 0.100, 1.000), max_depth (1, 2, 4, 8, 16, 30), and subsample (0.4, 0.5, 0.6, 0.7). Parameters gamma, colsample_bytree, and min_child_weight were set to 0, 0.8, and 1, respectively. The best-performing hyperparameter combination identified through grid search was then used to train the final model. TabPFN was performed using the TabPFNRegressor class from the TabPFN Python library (v2.0.3)[48].

### Evaluating the predictions

The coefficient of determination (*R^2^*) was used as the metric to evaluate our models. *R^2^* was also used in other studies that utilised microbiome data to predict environmental parameters[21, 49–51], which makes our results comparable to other efforts. The *R^2^* was calculated using the formula:

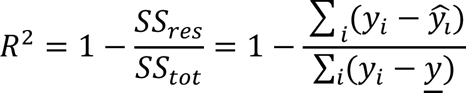

where *SS_res_* and *SS_tot_* are the residual and total sums of squares, respectively. *y_i_* is the actual value of the parameter, 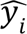 is the predicted value of the parameter for the same samples, and <u>*y*</u> is the mean of the actual values for the parameter.

When comparing machine learning to correlating the relative abundances of ASVs with microscopy, for the latter approach, we used the *R^2^* for linear regression between the microscopy and metabarcoding data. In other words, for the correlation, 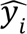 corresponds to the value predicted by the linear fit found with linear regression.

The classifiers, i.e., the models predicting presence/absence of phytoplankton and zooplankton taxa, were evaluated using F1-score. F1-score is a harmonic mean of precision and recall, and was calculated using the MLmetrics (v. 1.1.3) R package. The formula for F1-score is:

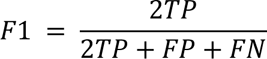

Where TP is the number of true positives, FP is the number of false positives, and FN is the number of false negatives. We counted presence as a positive result and absence as a negative result.

### Predicting microscopy identifications of plankton taxa

To predict the presence/absence of phytoplankton and zooplankton genera as determined by microscopy analysis, we tried three different approaches. First, we used physicochemical data as features to predict the microscopy-determined presence or absence of each genus in each sample. Second, we used 16S rRNA gene or 18S rRNA gene ASVs to make the same predictions. Third, we used the presence/absence of the same taxa obtained from 18S rRNA gene metabarcoding to predict the microscopy-determined presence/absence. For training, we converted the microscopy count table into binary data, with all counts >0 changed to 1. The random forest trained on this data returned values between zero and one. For evaluation, we converted the predicted values >=0.5 to 1, and <0.5 to 0, thus treating the random forest predictor as a binary classifier (while keeping the uncertain predictions in the original output). In addition to predicting presence/absence, the same setup was used to predict microscopy counts (individuals per volume unit) of the plankton genera (presented in Supplementary Fig. S2), running Random forest in regression mode and using coefficient of determination (*R^2^*) for the evaluations.

### Training and predicting on different datasets

To test the transferability of the machine learning models, we trained and evaluated the models using data from different sampling years. Data collected during 2019–2020 was used to predict parameters from 2015–2017 using 16S rRNA gene ASV counts for training. The “different dataset” predictions were compared to predictions from cross-validation based on the 2015–2017 dataset or both datasets. The same parameters were used to train the Random Forest models as those used for the physicochemical data.

### Calculating Environmental Status indicators from predicted and measured values

We used the model trained with the “different dataset” approach to predict physicochemical parameters for the years 2015–2017, using the corresponding measured values as a reference.

Ecological Quality Ratios (EQRs) were then calculated for the sampled basins based on both predicted and measured values of dissolved inorganic phosphate, water transparency (Secchi depth), total nitrogen, and total phosphorus. The calculations were performed using the HELCOM Eutrophication Assessment Tool (HEAT), which was updated for the Third Holistic Assessment of the Baltic Sea by the Baltic Marine Environment Protection Commission (HELCOM) and is publicly available at https://github.com/ices-tools-prod/HEAT.

## Results

### Physicochemical properties of seawater can be accurately predicted from the microbiome with machine learning

Random Forest machine learning models were trained to predict physicochemical properties of seawater from amplicon sequence variant (ASV) metabarcoding data (normalised counts of ASVs; hereafter referred to as ASV counts), using 10-fold cross validation (Fig. 1A). Accuracy was assessed by comparing the true values to the predicted values for each model using the coefficient of determination (*R^2^*). Metabarcoding-based models accurately predicted physicochemical properties of seawater (Fig. 2A & B). 16S rRNA gene data outperformed 18S rRNA gene data in predictive accuracy across all parameters, except chlorophyll-a (Chl-a). For the models based on 16S rRNA gene, the median *R^2^* was 0.844, illustrated by the prediction of nitrate (Fig. 2C). Salinity and alkalinity were predicted with the greatest accuracy (*R^2^* = 0.981 and 0.979, respectively), whereas ammonium (NH_4+_) and chlorophyll-a had the poorest (*R^2^*= 0.491 and 0.249). Overall, physicochemical parameters ranked similarly in terms of predictive performance across 16S and 18S rRNA gene-based models, although 16S rRNA gene-based models had higher overall accuracy.

**Fig. 2.**
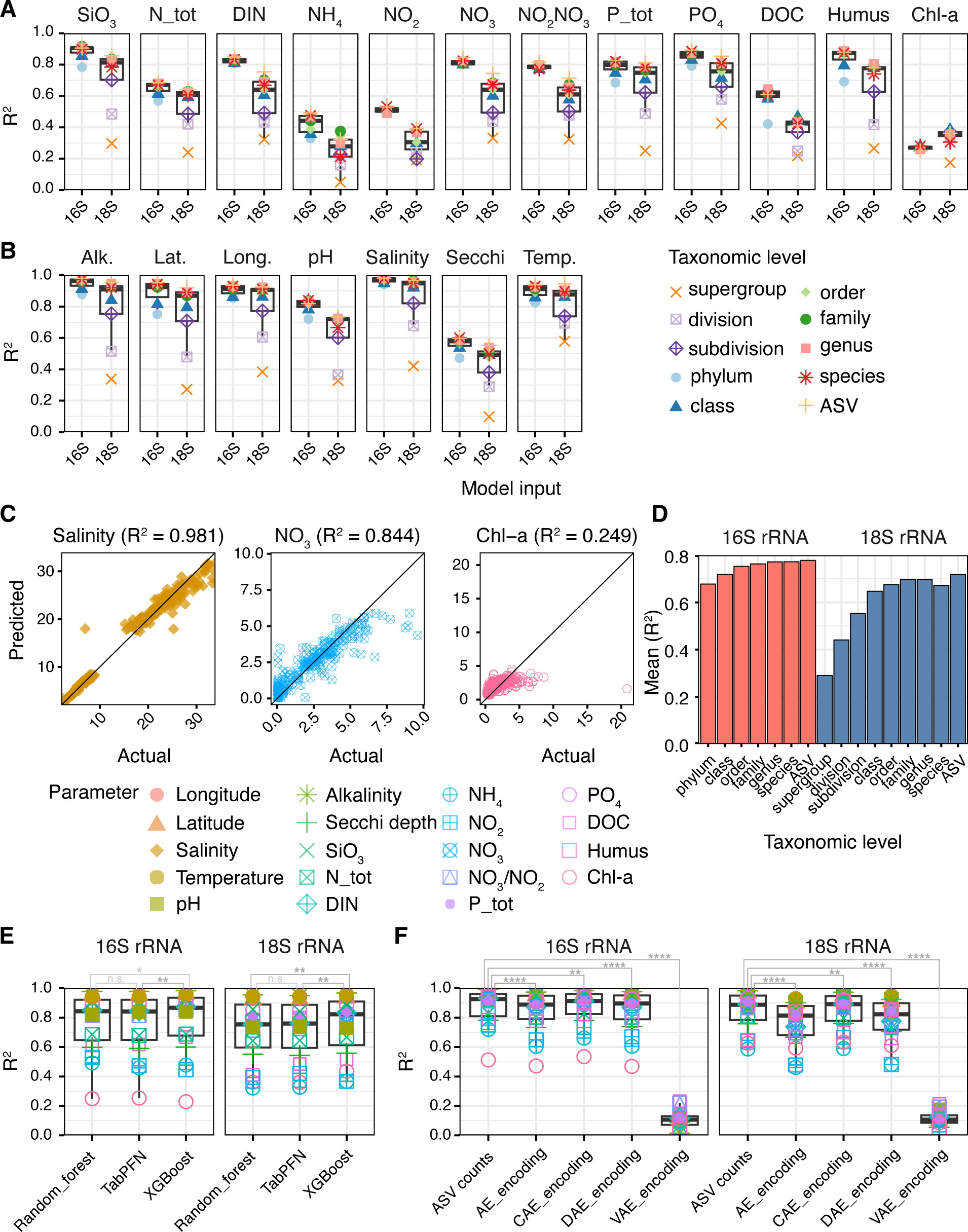
Performance of machine learning models for predicting physicochemical parameters from metabarcoding data. In all plots, performance is presented as coefficient of determination (*R*^2^) between observed and predicted values. **(A)** Predictive performance of Random Forest models for nutrient and biological parameters using metabarcoding data aggregated at different taxonomic levels. **(B)** Predictive performance of Random Forest models for physical parameters using metabarcoding data aggregated at different taxonomic levels. **(C)** Scatter plots of actual versus predicted data for best (Salinity), median (NO_3_), and worst (Chl-a) performing Random Forest models using 16S rRNA gene ASV data. **(D)** Mean model performance at different taxonomic levels across all environmental parameters. **(E)** Predictive performance of three machine learning algorithms: Random Forest, XGboost, and TabPFN, for predicting all physicochemical data from ASV data. **(F)** Predictive performance of Random Forest models with deep encoding architectures: shallow autoencoder (AE), convolutional autoencoder (CAE), deep autoencoder (DAE), variational autoencoder (VAE), for predicting all physicochemical data from ASV relative abundance. For each type of autoencoder, different architectures were tested, and only the best performing is shown. Asterisks indicate significance: \**P* < 0.05, \*\**P* < 0.01, \*\*\**P* < 0.001.

We tested whether aggregation of data at higher taxonomic levels would improve prediction performance compared to using ASVs. Models trained with ASVs as input features had greater predictive accuracy than models trained with data aggregated at higher taxonomic levels for both 16S rRNA gene and 18S rRNA gene data (Fig. 2D). For models using 16S rRNA gene data, prediction performance decreased progressively with each step of taxonomic aggregation (ASV to phylum). In contrast, for models using 18S rRNA gene data, aggregation at the family level resulted in the next best performance, surpassed only by the ASV-based predictions. As the ASV level performed best for both marker genes, ASV data was used as input for all subsequent analyses.

Using ASV-level metabarcoding data as input, we compared two alternative machine learning approaches, XGBoost and TabPFN, to Random Forest. All three approaches demonstrated strong predictive capabilities, underscoring the efficacy of metabarcoding data for predicting physicochemical properties of seawater. XGBoost performed significantly better than Random Forest and TabPFN for both 16S and 18S rRNA gene data, whereas Random Forest and TabPFN were comparable (Fig. 2E). Random Forest, however, required significantly less computational resources with comparable overall performance to XGBoost (mean *R^2^* 0.786 for XGBoost versus 0.778 for Random forest with 16S rRNA ASV data). We therefore selected Random Forest for the remaining analyses.

Next, we transformed the metabarcoding data into lower-dimensional embeddings using deep representation learning techniques prior to training the Random Forests [25], to assess whether this would improve prediction performance. Out of the embedding methods, the convolutional autoencoder (CAE) yielded the best results (Fig. 2F). However, training on raw ASV-level data outperformed all deep encoding embeddings (Fig. 2F), and was therefore used in the subsequent analyses.

### Metabarcoding data outperforms microscopy-based monitoring data for predicting abiotic parameters

The monitoring of phytoplankton and zooplankton with microscopy is an integral component of many marine monitoring programs, and is used to assess ecosystem status. We therefore compared the effectiveness of microscopy to the best performing metabarcoding data (16S ASV data) for predicting physicochemical parameters (Fig. 1B). Random Forest models trained with metabarcoding data performed consistently better than models trained on microscopy counts of both phytoplankton and zooplankton (Fig. 3). The only exception was chlorophyll-a, the parameter most poorly predicted with metabarcoding, where predictions worked best, though still poorly, with phytoplankton microscopy data (Fig. 3A).

**Fig. 3.**
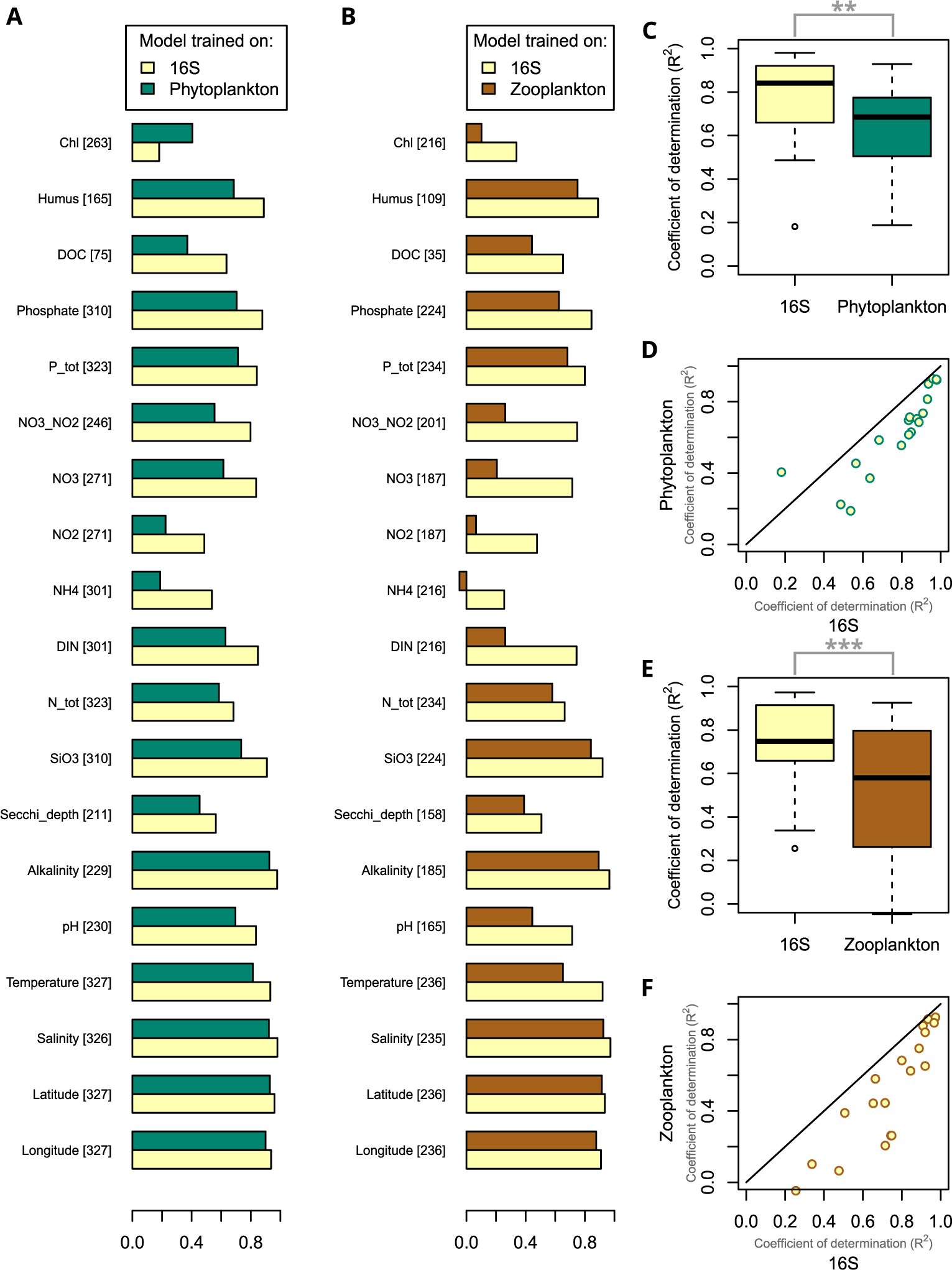
Performance of physicochemical parameter predictions using metabarcoding versus microscopy data. Machine-learning (Random Forest) trained on either 16S rRNA gene ASV counts or microscopy counts of **(A, C, D)** phytoplankton or **(B, E, F)** zooplankton. The microscopy-counted taxa were identified to either the genus or species level. Numbers in brackets in **A** and **B** indicate the number of samples included; only samples with both metabarcoding and microscopy data of either phytoplankton or zooplankton were included. Asterisks indicate significance: * *P* < 0.05, ** *P* < 0.01, *** *P* < 0.001.

### Metabarcoding can predict the presence of both phytoplankton and zooplankton taxa

Our results show that metabarcoding data accurately predicts physicochemical parameters of seawater (Figs. 2–3). We next assessed whether it can also predict biotic features (Fig. 1B). In this case, we focused on predicting the presence of specific phytoplankton and zooplankton genera, and compared training on either metabarcoding or physicochemical data since the latter is often the basis for generating species distribution models of plankton [52–54]. For each eukaryotic plankton genus detected with both microscopy and metabarcoding (according to the taxonomic annotation of the sequences) we predicted its presence or absence in the microscopy data with Random Forest models trained on (1) physicochemical data, (2) 16S rRNA ASV data, (3) 18S rRNA ASV data, and compared these predictions to (4) using the presence/absence of the genus in the 18S rRNA gene data (Fig. 4). For both phytoplankton and zooplankton, machine learning predictions with ASV counts had accuracies on par with those based on physicochemical data, and 16S and 18S ASV data performed equally well (Fig. 4C and 4I). As expected, the accuracy of the predictions increased with the number of samples in which the genus was identified with microscopy (Fig. 4A-B, D, G-H, J). Machine learning using ASV counts consistently outperformed using the presence/absence of the genus in the 18S rRNA gene data (Fig. 4F and L). This was most pronounced for zooplankton, where median F1-score with machine learning were ∼0.96 compared to ∼0.56 using the relative abundance of the corresponding genus in the metabarcoding data (Fig. 4I). In addition to predicting presence/absence, we predicted the counts obtained with microscopy of each taxon. Performance was considerably worse than predicting presence/absence, but the overall trends were the same (Supplementary Fig 2).

**Fig. 4.**
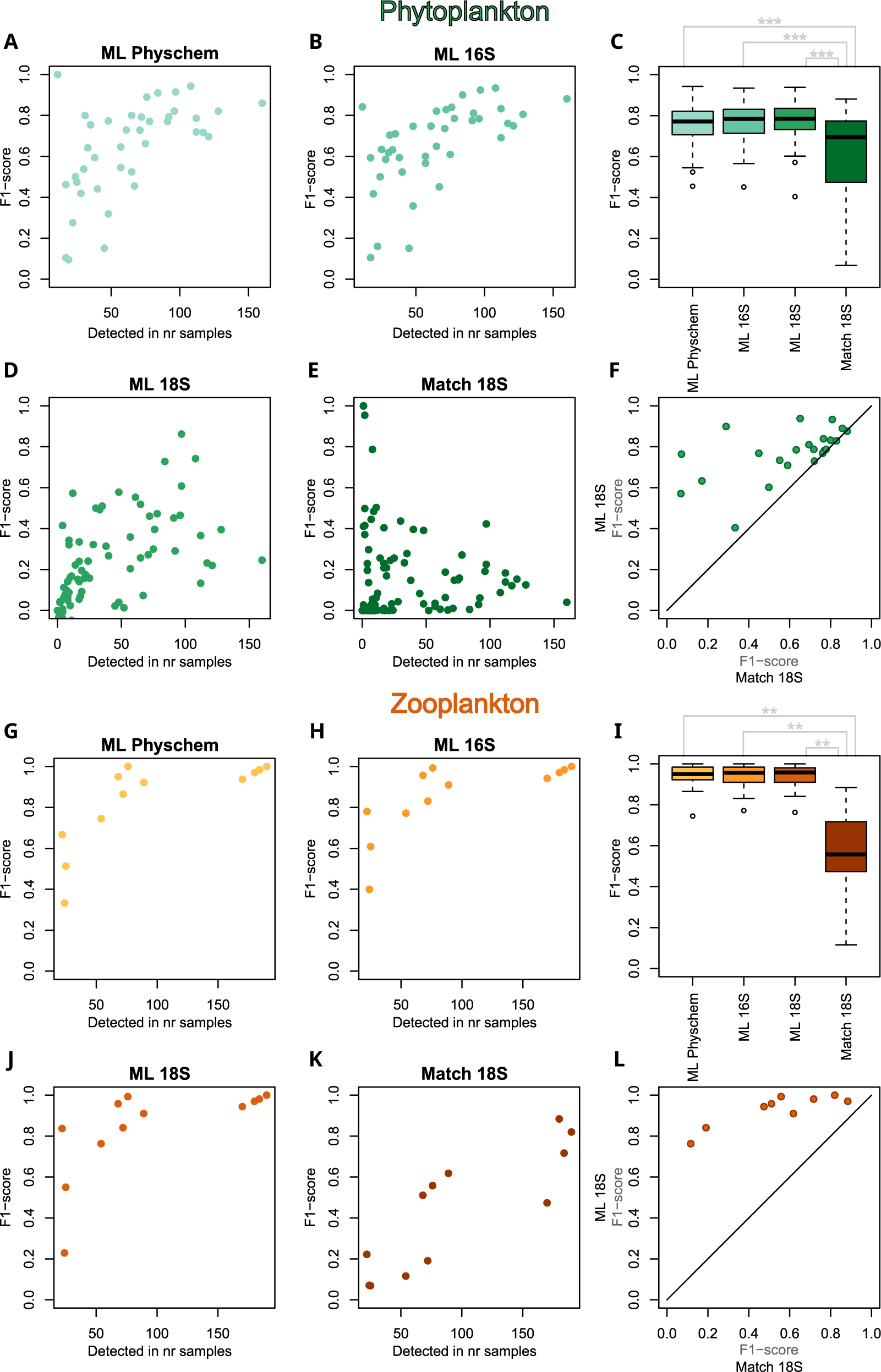
Performance of methods to predict the presence of phytoplankton (A-F) and zooplankton (G-L) genera. The presence or absence of each plankton genus in individual samples was predicted using Random Forest models trained on physicochemical variables (ML Physchem), 16S rRNA gene ASV count profiles (ML 16S), or 18S rRNA gene ASV count profiles (ML 18S). In addition, genus presence/absence was inferred directly from 18S rRNA gene data (Match 18S). Only genera detected with microscopy and metabarcoding (according to the taxonomic annotation of the sequences) in at least one sample each were included. In **C, F, I** and **L**, only genera detected with microscopy in >50 samples were included. “Detected in nr of samples” refers to the number of samples in which a given genus was identified by microscopy. For phytoplankton, 328 samples and for zooplankton, 236 samples were analysed. Asterisks indicate significance: * *P* < 0.05, ** *P* < 0.01, *** *P* < 0.001.

### Interannual predictions across datasets

We evaluated the temporal transferability of our machine learning models by training them on data from one time period and evaluating them on data from a different, non-overlapping time period (Fig. 1C). Specifically, we compared three strategies for predicting physiochemical data: (1) training on a different dataset (2019–2020) to the test data (2015–2017), (2) training with the same dataset as the test data (2015–2017), and (3) training using both datasets (2015–2017 and 2019–2020) for predicting the test data. We chose to use the 2015–2017 dataset as the test data, due to its narrower geographic coverage relative to the 2019–2020 data, thus avoiding extrapolation outside the geographic environmental gradients of the training data (Fig. 1C).

The performance of the interannual predictions (“different dataset”) compared to using the “same dataset” or “both datasets” models varied between parameters (Fig. 5A). In general, using the same dataset worked better than using a different dataset, and using both datasets worked best (Fig. 5B). However, for the six parameters best predicted with the “same dataset” model (*R*^2^ > 0.8), the interannual predictions performed comparably (Fig. 5C; *P* = 0.56).

**Fig. 5.**
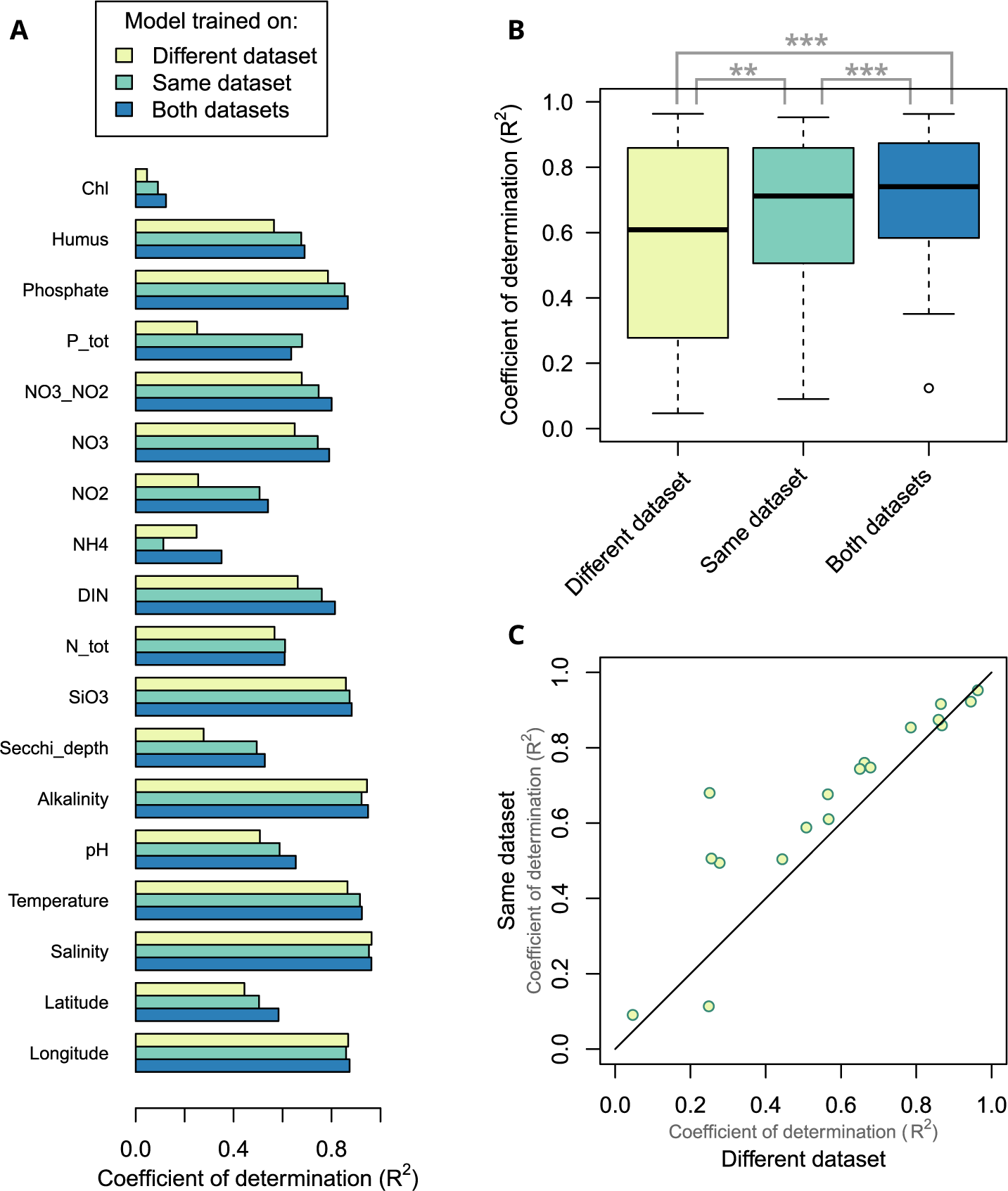
Transferability of machine learning models. Performance of Random Forest models for predicting physicochemical parameters from metabarcoding data using different datasets. Parameters were predicted for the 2015–2017 dataset with models trained on(1) the 2019–2020 dataset (“different dataset”), (2) the 2015–2017 dataset (“same dataset”), or (3) both 2015–2017 and 2019–2020 datasets (“both datasets”). Note that the “same dataset” and “both datasets” models used 10-fold cross-validation to ensure that the same samples were not used for predictions and training, as was also done in the other analyses. **(A)** coefficient of determination (*R*^2^) for each predicted parameter and training dataset. **(B)** Boxplots of *R*^2^ values for all parameters for the different training dataset (as shown in **A**). **(C)** Comparison of *R*^2^ values obtained from training on the “same dataset” or “different dataset”. Asterisks indicate significance: * *P* < 0.05, ** *P* < 0.01, *** *P* < 0.001.

We further assessed whether the poorer performance of some “different dataset” models reflect noisy or biased predictions. To this end, for all the predicted parameters, we obtained *R*^2^ values from linear regression between predicted and actual values, as a measure of variance, ignorant of bias. The linear regression-based *R*^2^ values did not significantly differ between the “different dataset” and “same dataset” models (*P* = 0.28), while the “both datasets” model was more precise than both other models (*P* < 0.01). This indicates that the poorer performance of the “different dataset” model for some parameters (e.g., total phosphorus; Supplementary Fig. S3B) is due to bias, i.e. systematic over or under estimation, rather than noise.

### Predicting indicators for good environmental status

To support ecological status assessments and inform management decisions in the Baltic Sea, the Baltic Marine Environment Protection Commission (HELCOM) uses a set of indicators across the themes: Biodiversity, Eutrophication, Maritime Activities, and Hazardous Substances [55]. For eutrophication, the Ecological Quality Ratio (EQR) values are used to assess deviation from reference conditions, where values close to one indicate good environmental status and values close to zero indicate poor ecological status [56].

We evaluated the potential of metabarcoding data to predict HELCOM eutrophication indicators using the “different dataset” model, and calculated the EQR based on both predicted and measured values of dissolved inorganic phosphate, water transparency (Secchi depth), total nitrogen, and total phosphorus. EQR estimates based on predicted physicochemical parameters matched those based on measured concentrations (*R*^2^ = 0.67, mean absolute error MAE = 0.058) (Fig. 6A). The assessment of eutrophication status based on measured data (Fig. 6B) is closely replicated by the predicted data (Fig. 6C).

**Fig. 6.**
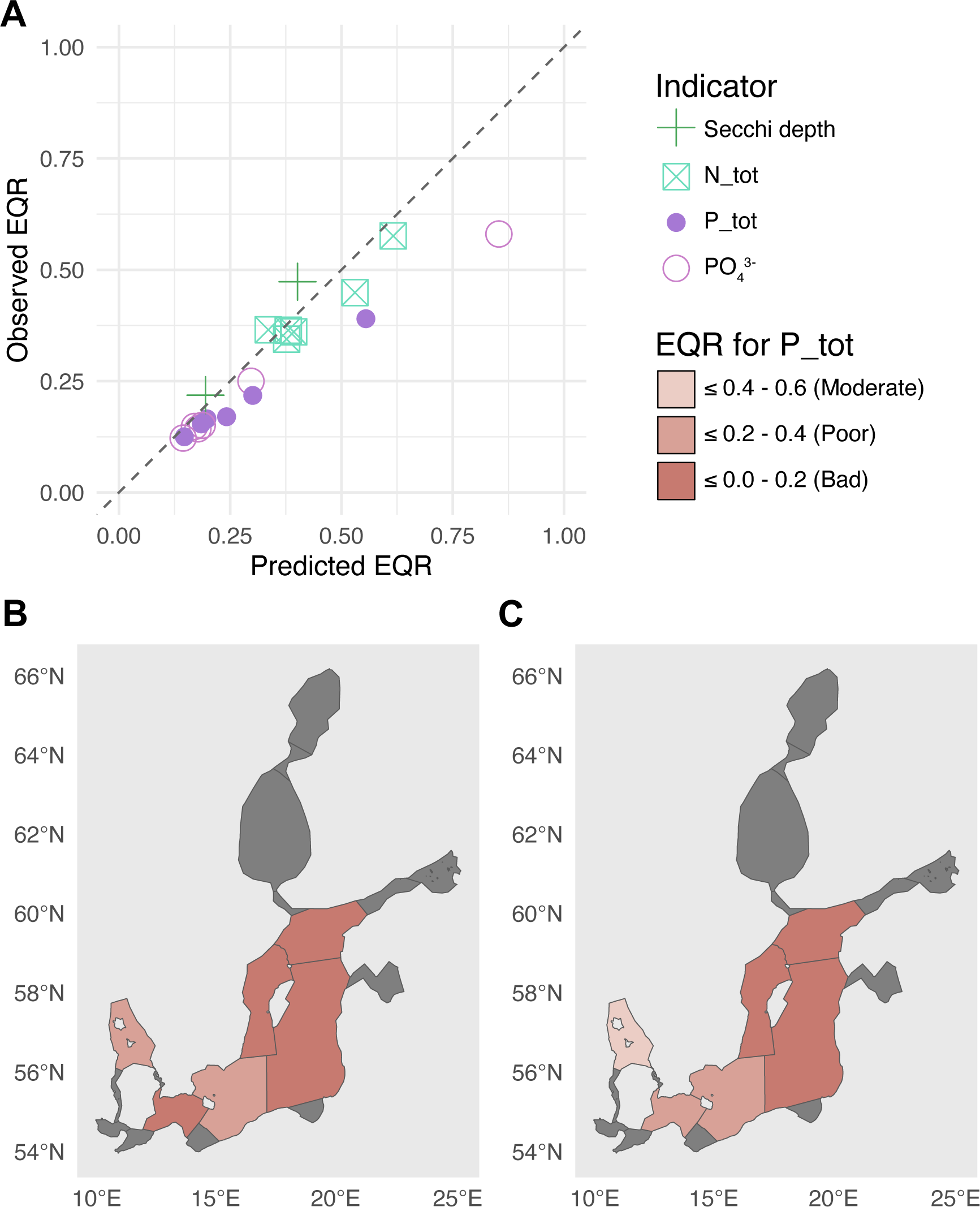
Calculating Ecological Quality Ratio (EQR) based on measured and predicted physicochemical parameters. Physicochemical parameters were predicted for the 2015–2017 dataset using the Random Forest algorithm trained with the 2019–2020 dataset (i.e., “different dataset”). **(A)** Observed and predicted EQR for water transparency (Secchi depth), total nitrogen (N_tot), total phosphorus (P_tot), and dissolved inorganic phosphate (PO_43-_), in different basins (Kattegat, Arkona Basin, Bornholm Basin, Eastern Gotland Basin, Western Gotland Basin, Northern Baltic Proper). EQR based on **(B)** measured values of P_tot, and **(C)** predicted values of P_tot.

## Discussion

Using metabarcoding data, we show that the taxonomic structure of microbial communities can be used to predict a range of both biotic and abiotic parameters in seawater. Using different machine learning approaches, physicochemical parameter predictions based on ASV-level 16S rRNA gene metabarcoding consistently outperformed those based on other feature types, showing that bacterial community structure holds a strong predictive power. Among machine learning models tested, the gradient boosting framework XGBoost achieved the highest overall accuracy but required substantially more computational resources than Random Forest. Random Forest models trained directly on ASV data outperformed those built using features extracted with autoencoders. Other studies have also found that feature selection does not necessarily improve the performance of ensemble tree methods for environmental prediction [51]. Considering both predictive performance and training time, we selected the Random Forest without feature extraction for predicting abiotic and biotic parameters from microbiome data. The generally high performance of all machine-learning architectures, however, demonstrates the overall robustness of metabarcoding data for predicting physicochemical parameters. These results highlight the potential of microbial community profiles to serve as biosensors for marine monitoring and to be integrated into predictive frameworks that enhance our understanding of ecosystem state and function.

Overall, model performance varied more between the predicted parameters than between the machine learning approaches used, indicating that data characteristics have a greater impact on model performance. The physicochemical parameters that were most accurately predicted – temperature, salinity, and alkalinity – are known to be important for structuring the marine microbiome. Both salinity [26, 36, 57] and temperature [13, 58] are key drivers of marine microbiome composition. The predictive power of these physical parameters likely reflects both their biological relevance and the consistency with which they are being measured. These parameters are easy to measure accurately with no difference in methodology between different sampling teams, instruments, or years, thus reducing potential analytical variability due to measurement limitations. In contrast, the physicochemical parameters that had the weakest predictive performance were chlorophyll-a and ammonia. The reason why predicting chlorophyll-a worked relatively poorly is not fully understood, though ammonia is considered one of the most challenging nutrients to measure accurately in seawater analysis [59]. Furthermore, in our data, the majority of ammonia data values are low with a few outliers (Supplementary Fig. S1). Thus, both analytical variability and spread of data likely influence the model performance.

Model performance was greater when using ASV counts compared to higher taxonomic levels or microscopy data, highlighting the value of high-resolution microbial community information. This aligns with a previous study on lakes where Random Forest models based on operational taxonomic units (OTUs) outperformed those based on higher taxonomic levels [21]. Another study focusing on headwater streams reported better performance using order-level classification than OTUs, but this study used a different algorithmic approach (Discriminant Analysis of Principal Components) [24], unassessed here. In the case of 16S rRNA gene data, prediction accuracy monotonically increased with finer taxonomic levels. For 18S rRNA gene data, however, the best performance after ASVs was at the family level. This is potentially due to lower annotation rates below that level [36] and points to a limitation of 18S rRNA gene reference databases and/or the taxonomic resolution of the amplified region (V4). Future studies using long-read sequencing that enables the recovery of full-length 16S rRNA or 18S rRNA ASVs may have the potential to further improve prediction performance.

Microscopy-based identifications of phytoplankton and zooplankton genera were better predicted with full ASV profiles than with the direct annotation of amplicon sequences with taxonomic databases. As the difference was strongest for zooplankton, we interpret that this reflects that the sample volume used for metabarcoding (<1 L) is too small to accurately quantify mesozooplankton taxa that are present in low concentrations. In comparison, cubic meters of water are sampled with zooplankton nets for microscopy counting. Given this, it is remarkable that machine learning using microbiome data from the small sample volumes can so accurately predict the presence/absence of zooplankton taxa.

Our results show that models trained on data from different years (“different dataset” models) can achieve similar accuracy to models trained on the same dataset for parameters that are well predicted by the latter. However, for other parameters, “different dataset” predictions were systematically worse. In general, these models were comparably precise as the “same dataset” models but less accurate due to biased predictions. For a few parameters, the “different dataset” models performed unusually worse than the other models. In these cases, the “different dataset” models gave similar predictions irrespective of actual values (Secchi depth and nitrite, Supplementary Fig. S3A and S3C) or underestimated the actual values (total phosphorus, Supplementary Fig. S3B). The poorer “different dataset” predictions may arise from differences in physicochemical measurements between the sampling years. Models may also be less transferable between datasets due to contextual differences between sampling years. Importantly, the “both dataset” model was both more accurate and precise than the “same dataset” model, showing that increasing the volume and diversity of training data improves model performance.

Using metabarcoding of the microbiome as a biosensor enables a multitude of information to be retrieved from a single sample, using one preparation and analysis pipeline. Given its strong predictive power for a range of physicochemical and biological variables, this approach could most likely be extended to infer new targets such as pollutants or toxins [60]. We explored its potential as an indicator of the Baltic Sea ecosystem health. Given that EQR ranges from 0 to 1, the level of predictive accuracy achieved (MAE = 0.058) suggests sufficient resolution to support ecological classification or to flag potential eutrophication hotspots. It could also be used to screen for locations with potential changes in environmental status, which could then be confirmed with more direct measurements. This approach offers a cost-efficient and scalable complement to traditional monitoring strategies, which often require separate sampling protocols and laboratory analyses for each parameter. Our findings are in line with earlier work demonstrating that microbiome-based machine learning models can reliably predict biotic indices to assess environmental impacts, for example, in marine aquaculture [31]. This supports the use of microbial profiles as indicators of ecosystem health.

In conclusion, this study highlights the ability of 16S and 18S rRNA gene metabarcoding to serve as biosensors for marine environmental monitoring. Using machine learning, we accurately predicted a range of biotic and abiotic variables, including key HELCOM indicators. Bacterial ASV data consistently yielded the highest performance across a broad range of parameters. By comparing the predictability of different variables, we map the strengths and challenges of metabarcoding-based machine learning. Our findings show that microbial community data, combined with robust models and diverse training datasets, can inform ecological assessments across time and space, offering a scalable tool for environmental monitoring.

## Data availability statement

The datasets analysed were previously published [35, 36] and are available as raw data in the European Nucleotide Archive (ENA) under accessions PRJEB5529 and PRJEB84926 and as processed data in the Swedish ASV portal at https://asv-portal.biodiversitydata.se. Code and processed metabarcoding and environmental data are available at https://github.com/EnvGen/EnvPredict.

## Authorship

Conceptualisation - all authors; Data Curation - all authors; Methodology - all authors; Formal analysis - EB, KG, KTJ, AFA; Visualisation - EB, KG, KTJ, AFA; Writing (original draft) - EB, KG, KTJ, AFA; Writing (review & editing) - all authors.

## Acknowledgements

The computations were enabled by resources provided by the National Academic Infrastructure for Supercomputing in Sweden (NAISS), partially funded by the Swedish Research Council through grant agreement no. 2022-06725.

