## Supplementary figures for "The marine microbiome can accurately predict its chemical and biological environment"

Supplementary Figures S1-S3.

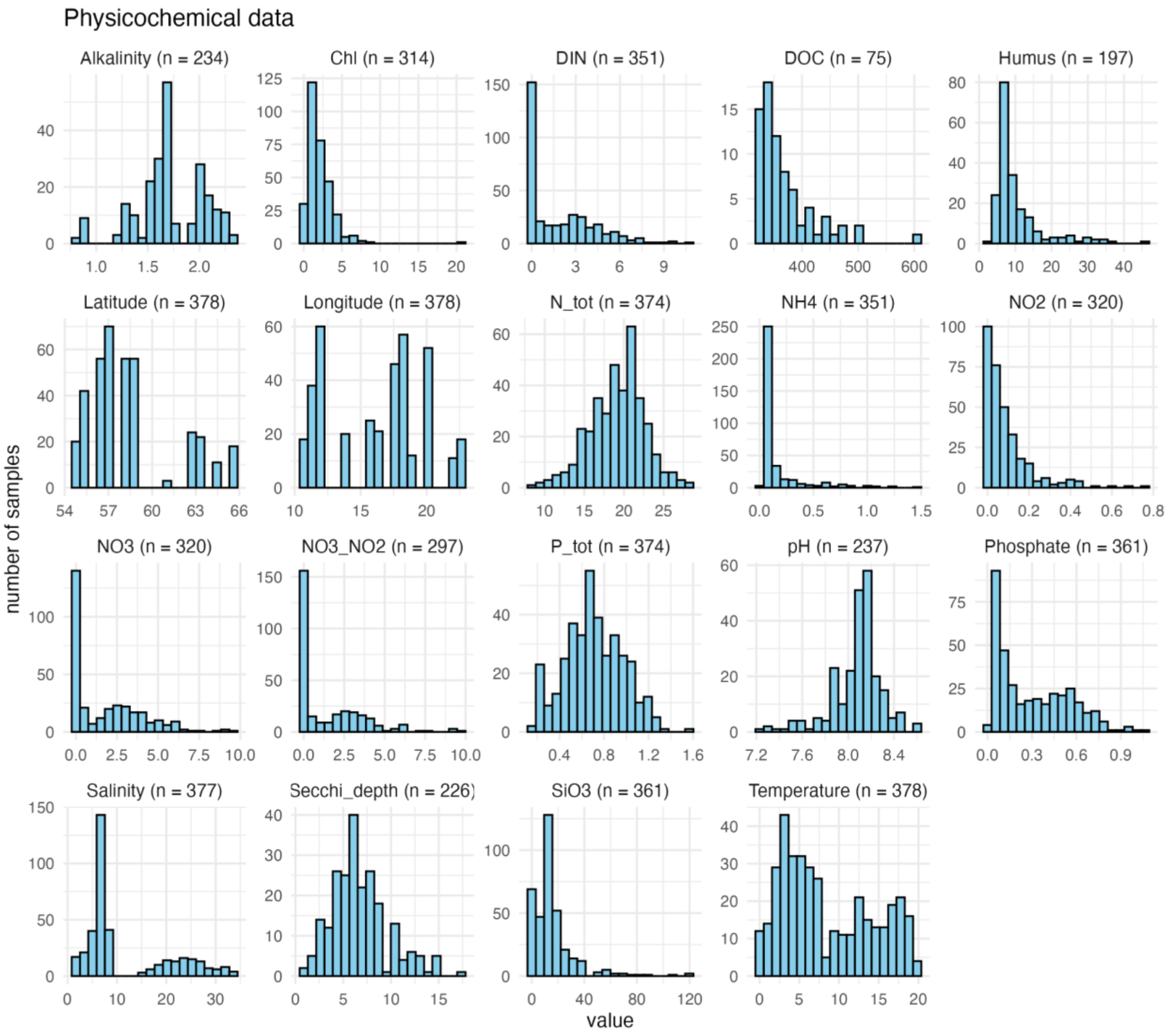


**Supplementary Fig. S1. Histograms showing data distribution of all physicochemical parameters predicted in this study.** Numbers in parenthesis denote the total number of measured values in the dataset.

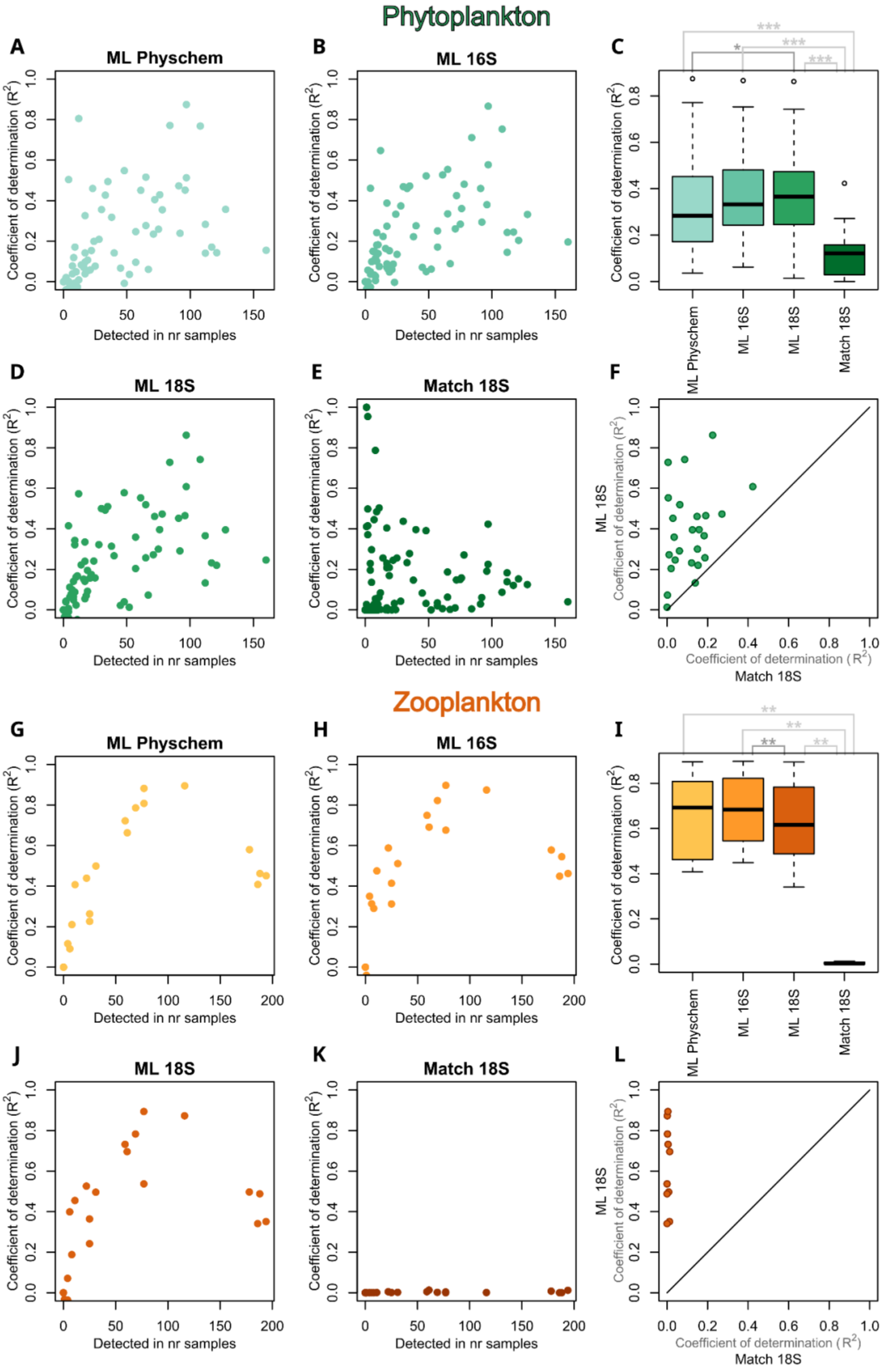

Phytoplankton
A
ML Physchem
B
ML 16S
C
D
ML 18S
E
Match 18S
F
Zooplankton
G
ML Physchem
H
ML 16S
I
J
ML 18S
K
Match 18S
L
Coefficient of determination (R2)
Detected in nr samples
ML Physchem
ML 16S
ML 18S
Match 18S
ML 18S
Coefficient of determination (R2)
Match 18S
***
*
**

**Supplementary Fig. S2. Performance of methods to predict counts of phytoplankton (A-F) and zooplankton (G-L) genera.** The counts (individuals per sampled water volume) of each plankton genus in individual samples was predicted using Random Forest models trained on physicochemical variables (ML Physchem), 16S rRNA gene ASV count profiles (ML 16S), or 18S rRNA gene ASV count profiles (ML 18S). In addition, the counts were inferred directly from the relative abundances of the genus in the 18S rRNA gene data (Match 18S). Only genera detected with microscopy and metabarcoding (according to the taxonomic annotation of the sequences) in at least one sample each were included. In **C, F, I** and **L**, only genera detected with microscopy in >50 samples were included. “Detected in nr of samples” refers to the number of samples in which a given genus was identified by microscopy. For phytoplankton, 328 samples and for zooplankton, 236 samples were analysed. The $R^2$ for the ‘Match 18S’ option was calculated based on linear regression (agnostic to the slope) since relative sequence abundance cannot readily be translated into number of organisms per volume, while for the machine learning, coefficient of determination relative to the identity line between the predictions and actual data was used. Asterisks indicate significance: * $P < 0.05$, ** $P < 0.01$, *** $P < 0.001$.

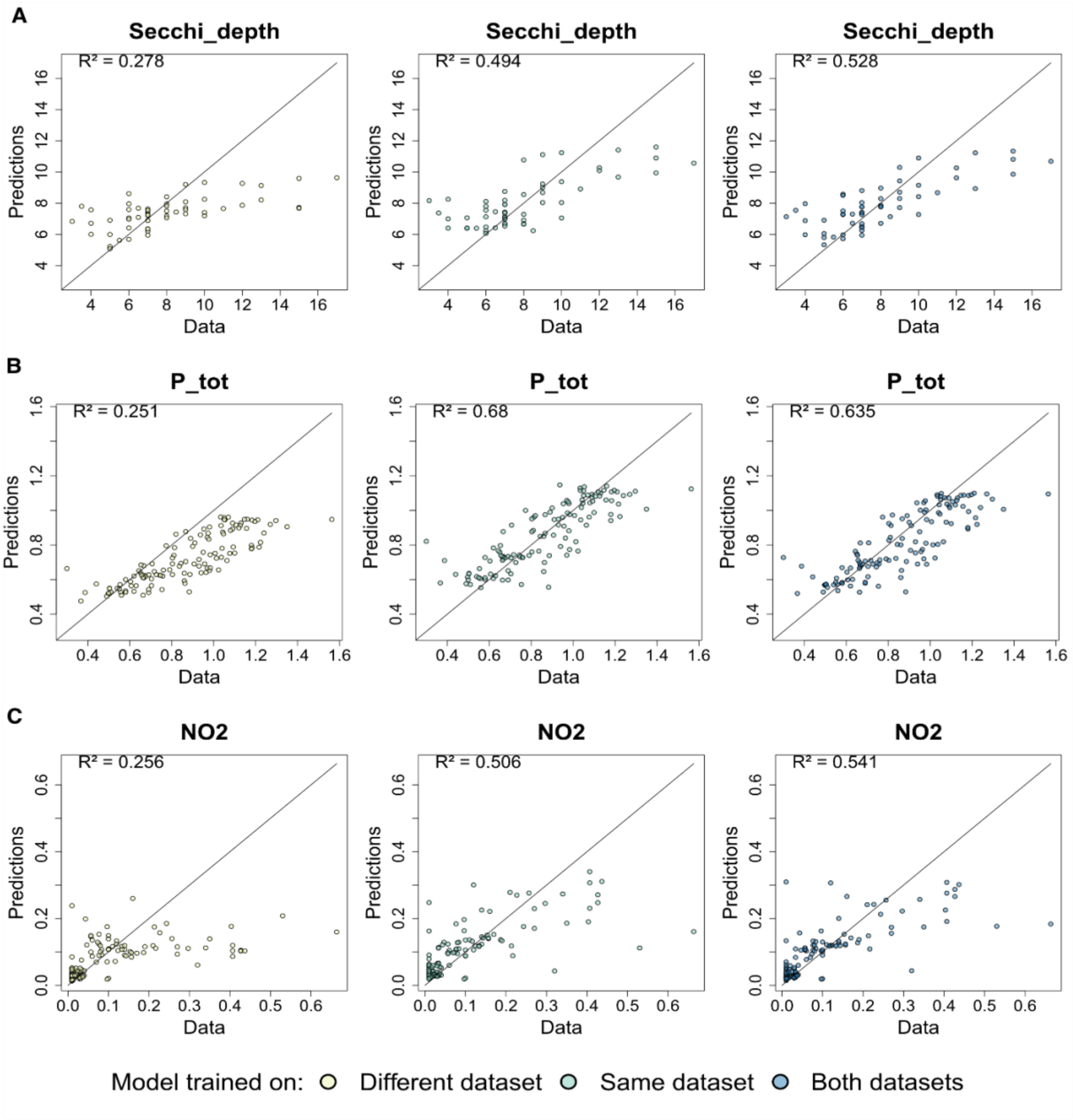


**Supplementary Fig. S3.** Predicted vs actual values of selected parameters with unusual performance patterns, comparing training on a different dataset (left panels), the same dataset (middle panels), and both datasets (right panels). **(A)** Secchi depth **(B)** Total phosphorus **(C)** Nitrite.
